# FusB energises import across the outer membrane through direct interaction with its ferredoxin substrate

**DOI:** 10.1101/749960

**Authors:** Marta Wojnowska, Daniel Walker

## Abstract

Phytopathogenic *Pectobacterium* spp. import ferredoxin into the periplasm for proteolytic processing and iron release via the ferredoxin uptake system. Although the ferredoxin receptor FusA and the processing protease, FusC, have been identified, the mechanistic basis of ferredoxin import is poorly understood. In this work we demonstrate that protein translocation across the outer membrane is dependent on the TonB-like protein FusB. In contrast to the loss of FusC, loss of FusB or FusA abolishes ferredoxin transport to the periplasm, demonstrating that FusA and FusB work in concert to transport ferredoxin across the outer membrane. In addition to interaction with the TonB-box region of FusA, FusB also forms a complex with the ferredoxin substrate, with complex formation required for substrate transport. These data suggest that ferredoxin transport requires energy transduction from the cytoplasmic membrane via FusB for both removal of the FusA plug domain and for substrate translocation through the FusA barrel.

## Introduction

Gram-negative bacteria have evolved a number of strategies for the acquisition of iron and other nutrients in which TonB-dependent transporters (TBDTs) play a central role^1^. In the case of siderophore-mediated iron acquisition the iron-siderophore complex is imported into the cell, captured by a siderophore-specific periplasmic binding protein and delivered to an ABC-transporter for import to the cytoplasm^2^. For iron acquisition from large host proteins such as transferrin, the iron-containing protein is captured at the cell surface through TBDT binding and the iron stripped at the cell surface and subsequently transported through the lumen of the TBDT^3^. In addition to the outer membrane receptor, whose lumen constitutes the translocation route, TBDT-mediated transport requires a complex of three proteins anchored in the inner membrane: TonB, ExbB and ExbD^4,5^. The ExbBD-TonB complex allows the entry of the nutrient by removal of a force-labile portion of the plug domain, which obstructs the receptor lumen^6^. ExbB and ExbD are related to the flagellar motor proteins and harness proton motive force to energise the transport process.

In addition to the uptake of iron siderophores and other metal chelating compounds such as vitamin B_12_, TBDTs also transport complex carbohydrates and simple sugars^7^. A recent study has also described the role of a TonB-dependent receptor in protein export, suggesting that TonB-dependent receptors are highly adaptable to the transport of diverse substrates across the OM^8^. The flexibility in the range of substrates that are amenable to transport by TBDTs is exploited by protein antibiotics such as colicins and pyocins that use TBDTs as their primary cell surface receptor and translocator^9^. As with the uptake of nutrients, translocation of colicins and pyocins via TBDTs is PMF-dependent, although in these cases the periplasm-spanning protein TonB is required both to remove the force labile region of the TBDT-plug domain and to subsequently energise protein translocation across the OM^10^. Protein translocation occurs by direct interaction with the an N-terminal intrinsically unstructured region of the toxin that, similar to the TBDTs, carries a TonB-binding motif^10^.

We recently demonstrated that TBDT-mediated iron acquisition from the iron-sulphur cluster containing protein ferredoxin represents an unprecedented example of protein translocation into the bacterial cell for nutrient acquision^11^. Ferredoxin binding at the cell surface is mediated by the TBDT FusA and, following transport of intact ferredoxin into the periplasm, the substrate is subjected to proteolytic processing by the M16 protease FusC^11,12^. Cleavage by FusC results in release of the iron-sulphur cluster and is required for effective iron acquisition from ferredoxin by *Pectobacterium*. Together with the genes encoding FusA and FusC, the Fus operon contains two additional genes, with *fusB* encoding a TonB-homologue and *fusD* a putative ABC transporter^12^. Interestingly, the M-type pectocins M1 and M2, which we have previously described, parasitise the ferredoxin uptake system through an N-terminal ferredoxin domain that is highly homologous to plant ferredoxins^13,14^.

More recently, the X-ray structure of FusC bound to ferredoxin has been reported, showing that substrate recognition occurs at a site distant from the active site^15^. Furthermore, only parts of the ferredoxin molecule are visible in the structure, implying that the bound substrate is largely unstructured. Based on these data, it was suggested that ferredoxin transport occurs by means of a Brownian ratchet mechanism in which FusC acts as a periplasmic anchor to facilitate translocation of ferredoxin across the OM via the lumen of FusA^12,15^. Similar mechanisms have been postulated to account for mitochondrial protein uptake, whereby cytoplasmically synthesised proteins are translocated via the TOM and TIM23 complexes^16,17^. As such, this would represent a hitherto unexpected evolutionary link between mitochondrial and plastid protein import and bacterial protein import via the Fus and other postulated protein uptake systems.

In this work we show that FusC does not facilitate ferredoxin import and that like the import of other TBDT substrates, ferredoxin uptake is PMF-dependent. Instead, we show that the TonB-homologue encoded within the *fus* operon, FusB, is required for ferredoxin import and the mechanism of ferredoxin import involves a direct interaction between FusB and the ferredoxin substrate. The direct interaction of the TonB-like protein with substrate is unprecedented and explains the requirement for the system-specific TonB-homologue in the Fus system. Our data also show that, in addition to the direct interaction with the substrate, FusB fulfils another role - similar to other TonB proteins - in interacting with the TonB-box of FusA for plug displacement. Since multiple genes encoding TonB-like proteins are commonly found in the genomes of Gram-negative bacteria this may be a common mechanism for the uptake of atypical substrates via TonB-dependent receptors.

## Results

### Ferredoxin import is independent of FusC but requires proton motive force

We previously showed that FusC is a highly specific protease that targets plant ferredoxin to release iron from this host protein in the periplasm of *Pectobacterium* spp^11^. However, it has also been suggested that FusC plays an additional role in iron acquisition through a direct involvement in ferredoxin transport across the outer membrane by means of a Brownian ratchet mechanism, specifically acting as a periplasmic anchor^15^. Our own previous work suggests that if FusC does play a role in ferredoxin import this role is not essential since the accumulation of Arabidopsis ferredoxin can still be observed in a *P. carotovorum* LMG2410 (*Pc*LMG2410) strain lacking FusC^11^. However, using Arabidopsis ferredoxin (Fer_Ara_) it is not possible to directly compare the rate and extent of ferredoxin uptake between wild-type and *ΔfusC P. carotovorum* since Fer_Ara_ is cleaved by FusC on import to the periplasm in the wild-type strain; hence, based on these data, we could not rule out a role for FusC in ferredoxin import.

To test the hypothesis that FusC facilitates translocation of ferredoxin to the periplasm we compared the uptake of potato ferredoxin (Fer_Pot_) in wild-type and *ΔfusC Pc*LMG2410. Fer_Pot_ was used as we had observed that, although similarly to Fer_Ara_, it can be transported into cells (Figure 1a), unlike Fer_Ara_ and spinach ferredoxin (Fer_Sp_), which both support robust growth of *Pc*LMG2410 under iron-limiting conditions^12,13^, it is not cleaved at an appreciable rate by FusC and so accumulates intracellularly in wild-type *Pc*LMG2410 (Figure 1b). To compare uptake of Fer_Pot_ in wild-type and *ΔfusC Pc*LMG2410, cells were grown under iron-limiting conditions through the addition of the iron chelator, 2,2’-bipyridine, and supplemented with Fer_Pot_. The amount of Fer_Pot_ in whole cell extracts and the media was determined by immunoblotting. Levels of Fer_Pot_ obtained from cell extracts increased at the same rate in whole cell extracts and rates of removal of ferredoxin from the media were similar (Figure 1c). These data show that FusC does not play a role in protein import and the role of FusC in iron acquisition is likely restricted to proteolytic processing of ferredoxin as we previously reported^11^.

**Figure 1.**
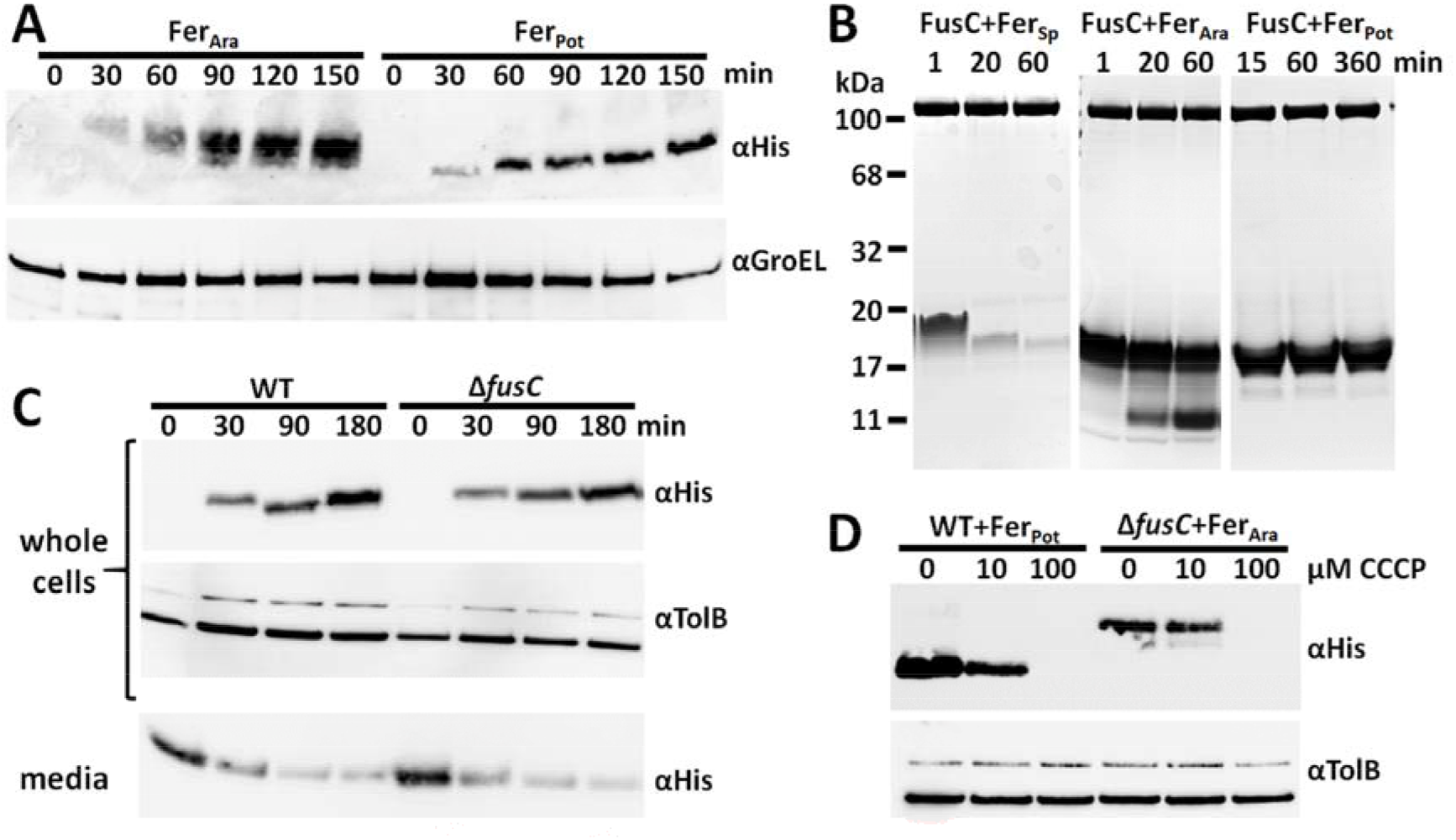
Import of ferredoxin is FusC-independent and requires proton motive force. (A) Uptake of Fer_Ara_ and Fer_Pot_ by *ΔfusC* cells over time determined by immunoblotting of whole cell extracts; GroEL serves as loading control. (B) FusC-mediated cleavage assays of plant ferredoxins. (C) Comparison of Fer_Pot_ uptake by wild-type (WT) and *ΔfusC* cells with TolB as the loading control; the bottom panel showing the concomitant depletion of ferredoxin from the media. (D) Ferredoxin import assays in the presence of protonophore CCCP; Fer_Pot_ was used as a reporter in wild type cells while Fer_Ara_ was used in *ΔfusC* cells.

To further probe the mechanism of ferredoxin uptake we tested the ability of *Pc*LMG2410 cells to accumulate Fer_Pot_ under iron-limiting conditions and in the presence of the uncoupling agent carbonyl cyanide m-chlorophenylhydrazone (CCCP), which dissipates the PMF through transport of protons across the cytoplasmic membrane^18^. The intracellular accumulation of Fer_Pot_ by *Pc*LMG2410 was markedly reduced in the presence of 10 μM CCCP, relative to cells grown in the absence of the uncoupling agent, and abolished at 100 μM CCCP (Figure 1d). Similar effects through the action of CCCP were observed on the intracellular accumulation of Arabidopsis ferredoxin by *ΔfusC* LMG2410 (Figure 1d). These data show that, analogously to other TBDT-mediated transport processes^10,19^, the import of ferredoxin is PMF-dependent and a Brownian ratchet mechanism is unlikely to play a key role in ferredoxin import in *Pectobacterium* spp.

### FusB mediates ferredoxin import into the periplasm

As we previously reported, in addition to *fusA* and *fusC,* the Fus operon carries additional genes that encode a TonB homologue, FusB, and an ABC transporter FusD^12^. Given the documented role of TonB in siderophore import in many bacterial species we supposed FusB may play a similar role in protein import, having perhaps evolved additional functionality required to mediate the passage of a large substrate through the lumen of the TBDT FusA. To test this hypothesis we created *ΔfusA* and *ΔfusB* strains in *Pc*LMG2410 and initially probed them using growth enhancement assays under iron-limiting conditions. As indicated by the loss of growth enhancement both on solid media (Figure 2a) and in liquid culture (Figure 2b), these two genes encode proteins which are essential for Fus-mediated iron acquisition. The possibility that deletion of either gene affected the expression or level of FusC, thus indirectly affecting the growth enhancement phenotype, was ruled out by immunoblotting whole cell extracts with anti-FusC antiserum (Figure S1). We further investigated the ability of the *ΔfusA* and *ΔfusB Pc*LMG2410 to import ferredoxin relative to wild-type and *ΔfusC* strains using Fer_Pot_, which cannot be cleaved by FusC. Consistent with the hypothesised role of FusB in protein import, and in contrast to wild-type and *ΔfusC* strains, we did not observe the intracellular accumulation of Fer_Pot_ in *ΔfusA* and *ΔfusB Pc*LMG2410 (Figure 2c). The ferredoxin import phenotype lost in the *ΔfusA* and *ΔfusB* strains was restored by plasmid-based complementation of *fusA* and *fusB,* respectively (Figure 2d). In these experiments the production of FusA and FusB is IPTG-inducible under the control of the T5 promoter, although in the case of FusB complementation, leaky expression in the absence of IPTG is sufficient to restore protein import.

**Figure 2.**
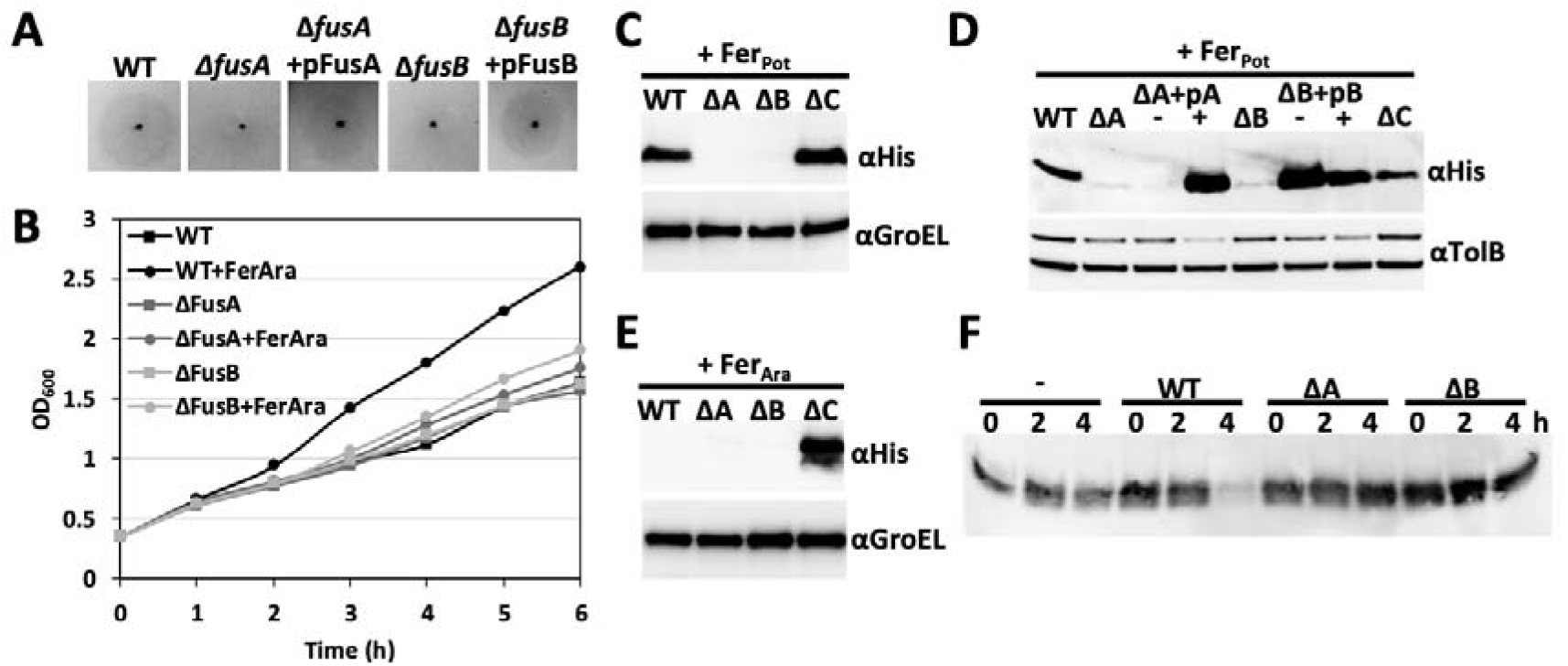
FusA and FusB are required for ferredoxin uptake. (A) Growth enhancement assay using spinach ferredoxin spotted onto soft agar overlay containing wild-type (WT), deletion strains and deletion strains recomplemented with the respective plasmid. (B) Growth curve comparing the rate of growth of wild-type and each deletion strain in the absence and presence of Arabidopsis ferredoxin. (D) Internalisation assay of Fer_Pot_; mid-log phase cells of each strain were supplemented with 2,2’-bipyridine and 1 μM ferredoxin, grown for 1 hour and whole cell extracts were probed with anti-His antiserum. (E) Internalisation assay of Fer_Pot_ including deletion strains recomplemented in *trans* using plasmids encoding the respective genes (“ΔA+pA” = Δ*fusA*+pFusA, “ΔB+pB” = Δ*fusB*+pFusB); prior to the addition of 2,2’-bipyridine and ferredoxin, the cultures of recomplemented strains were split in two and either supplemented with 0.5mM IPTG (“+”) or grown in the absence of inducer (“-”). (F) Internalisation assay using Fer_Ara_ (see D). (G) Depletion assay showing the gradual reduction of Arabidopsis ferredoxin level in the media derived from uninoculated (-), wild-type, *ΔfusA* (ΔA) and *ΔfusB* (ΔB) cell cultures.

To ensure that the abrogation of substrate import is not specific to Fer_Pot_ we also monitored the ability of the *ΔfusA* and *ΔfusB* strains to utilise Fer_Ara_. However, for this substrate instead of measuring intracellular ferredoxin accumulation, we determined loss of ferredoxin from the growth media, since accumulation of Fer_Ara_ is only observed in the LMG2410 *ΔfusC* strain (Figure 2e). Consistent with the internalisation assay and growth enhancement assays, ferredoxin content of the media decreased over time in the presence of wild type cells, but not in the presence of the *ΔfusA* and *ΔfusB* strains, indicating that both FusA and FusB are required for Fer_Ara_ uptake (Figure 2f).

### FusB directly interacts with the ‘TonB-box’ of FusA

Having determined that the TonB-like protein FusB is required for ferredoxin import we aimed to elucidate the mechanism of ferredoxin uptake. Analysis of the FusA sequence showed the presence of a putative TonB-box, DTILVRST, with a similar sequence to TonB-boxes from well-characterised *E. coli* TBDTs and other *Pc*LMG2410 TBDTs (Figure S2). The functional importance of this putative TonB-box region was demonstrated using ferredoxin import assay and plasmid-based complementation of the *ΔfusA* strain, which showed that proline substitutions within the putative TonB-box abolish internalisation of the FusA substrate ferredoxin (Figure S3). The similarity of the TonB-box of FusA to the TonB-boxes of other *Pc*LMG2410 and *E. coli* TBDTs suggests that FusA may interact with *Pc*LMG2410 TonB and not FusB. Although *Pc*LMG2410 has 6 genes that encode TonB-like proteins, we hypothesised that the protein that was most similar to *E. coli* TonB, which we refer to as *Pc*TonB, would fulfil the same function as this protein in servicing multiple TBDTs, with the primary function of dislocating the plug domain to enable substrate transport.

To determine if TonB and/or FusB interact directly with FusA we produced a construct consisting of the N-terminal region of FusA (residues 21 to 66), excluding the signal peptide region, fused to GFP (FusA_NTR_-GFP) and determined if this interacts with the isolated C-terminal domains of *Pc*TonB (TonB_CTD_) and FusB (FusB_CTD_) by ITC. Clear heats of binding were observed on titration of FusA_NTR_-GFP into FusB_CTD_, although the affinity of FusB_CTD_ for FusA_NTR_-GFP is weak (57 μM) (Figure 3a). No heats of binding were observed on titration of isolated GFP into FusB_CTD_, (Figure S4) showing that the C-terminal domain of FusB specifically interacts with the N-terminal region of FusA. Interestingly, similar heats of binding were observed on titration of FusA_NTR_-GFP into TonB_CTD_ (Figure S5), demonstrating that the N-terminal region of FusA can also interact with *Pc*TonB.

**Figure 3.**
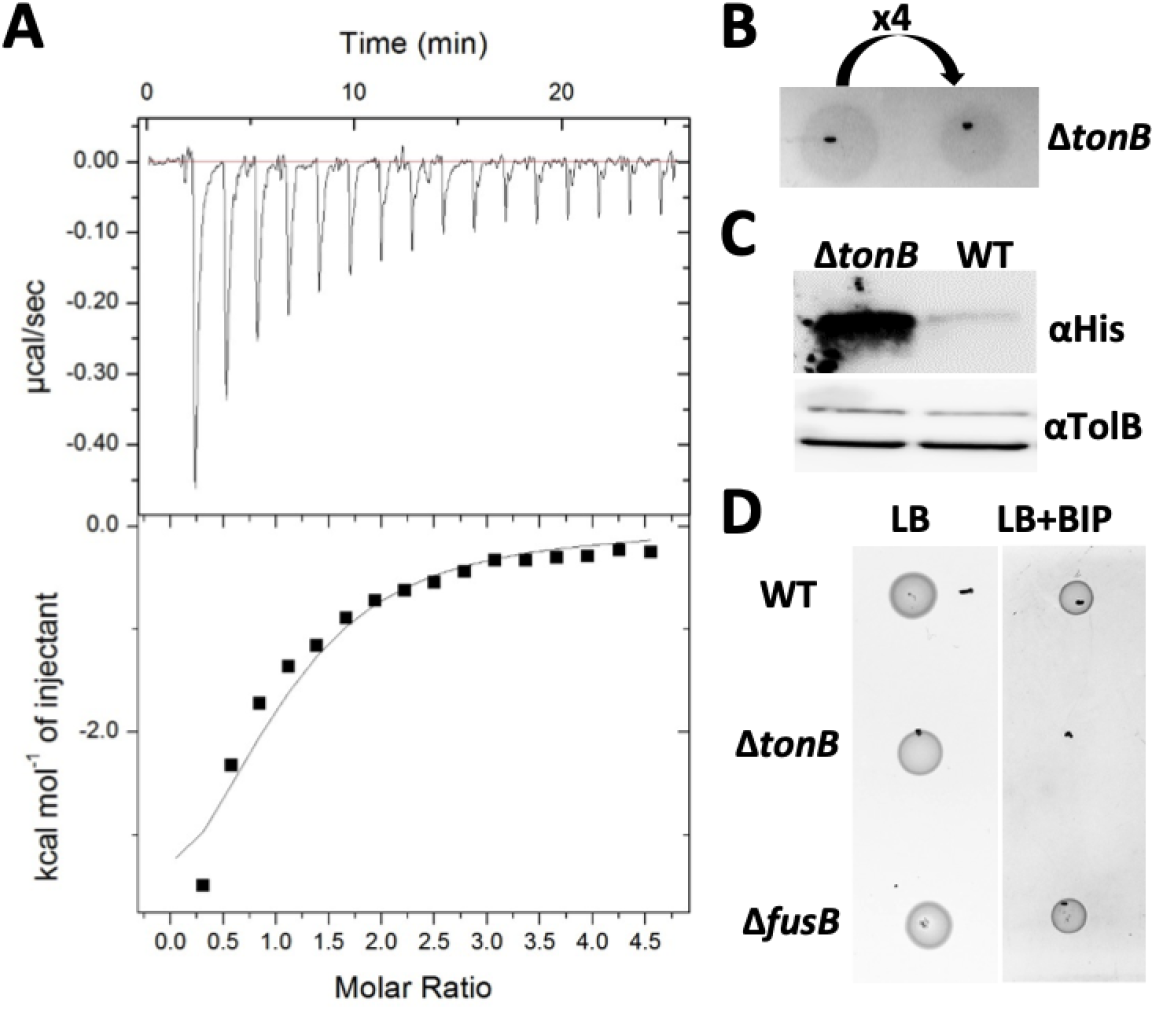
FusB interacts with the FusA ‘TonB-box’ and ExbB, but not *Pc*TonB, is required for ferredoxin uptake. (A) ITC binding isotherm of 1 mM FusA_NTR_-GFP titrated into 90 μM FusB_CTD_. The calculated K_d_ for the FusA_NTR_-GFP-FusB_CTD_ complex is 57(±12) μM (n=3). (B) Growth enhancement assay using spinach ferredoxin on soft agar overlay containing *ΔtonB* cells; x4 refers to dilution factor. (C) Uptake of potato ferredoxin by *ΔtonB* and wild-type cells over the course of 1 hour. (D) Growth assay of wild type (WT), *ΔtonB* and *ΔfusB* cells spotted onto LB agar (LB) or LB agar supplemented with 400μM 2,2’-bipyridine (LB+BIP).

To determine if the *Pc*TonB plays a role in ferredoxin uptake, we deleted the respective gene in *Pc*LMG2410 and tested the growth enhancement phenotype of this strain in the presence of Fer_Sp_. In contrast to deletion of *fusB,* the loss of *tonB* did not reduce growth enhancement phenotype (Figure 3b); in fact, the *ΔtonB* strain showed more prominent zones of growth enhancement in the presence of Fer_Sp_ relative to the wild-type strain and showed increased intracellular accumulation of Fer_Pot_ relative to wild-type *Pc*LMG2410 (Figure 3c). However, *Pc*LMG2410 *ΔtonB* exhibited poor growth in the presence of 2,2’-bipyridine relative to the wild-type strain (Figure 3d), with very faint growth observable after 24 hours. These data indicate that, although *Pc*TonB does play the expected generic role in iron uptake, this does not include iron acquisition from ferredoxin. Furthermore, despite the aforementioned observation that TonB interacts with FusA *in vitro* (and possibly *in vivo),* this interaction is not sufficient for ferredoxin uptake. Indeed there may be competition between FusB and TonB for complex formation with FusA, with only the FusB-FusA complex being productive with respect to ferredoxin uptake.

### FusB interacts directly with the ferredoxin substrate

The ability of FusB and *Pc*TonB to interact with FusA, but with only the former able to mediate ferredoxin uptake, suggests that FusB plays an additional role, which is essential for ferredoxin import. One possibility is that FusB directly interacts with the protein substrate after the initial binding of ferredoxin to FusA at the cell surface. To test this, we sought to determine if ferredoxin forms a complex with the isolated C-terminal domain of FusB by size exclusion chromatography (SEC). SEC of ferredoxin mixed with FusB_CTD_ monitored at 280 nm and 330 nm gave a peak indicating the presence of a species of higher molecular weight than FusB_CTD_ or Fer_Sp_ alone, providing evidence of complex formation (Figure 4a). In contrast, no complex formation was observed between ferredoxin and the purified C-terminal domain of TonB (TonB_CTD_) using SEC (Figure S6). We further investigated the formation of the FusB_CTD_-Fer_Sp_ complex by isothermal titration calorimetry, titrating Fer_SP_ into FusB_CTD_ (Figure 4b). These data show that FusB interacts directly with the ferredoxin substrate.

**Figure 4.**
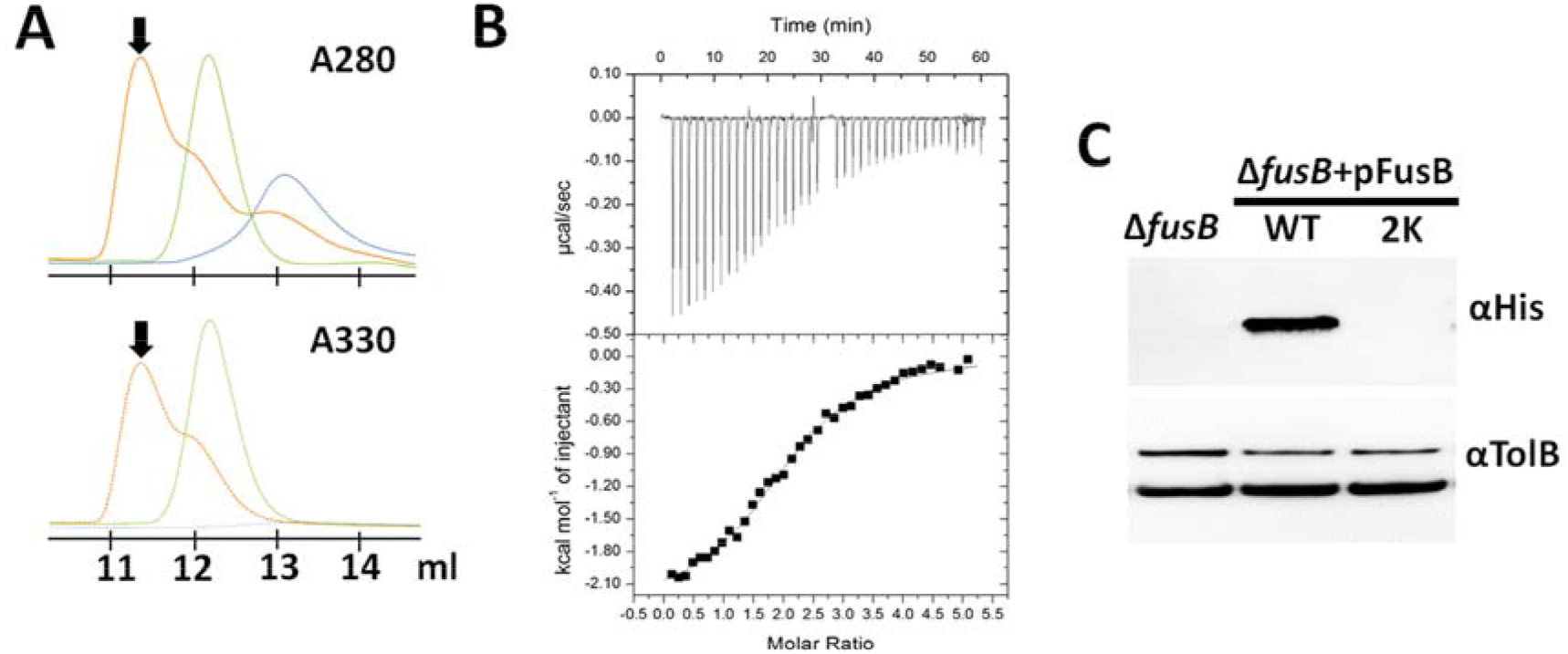
FusB interacts with ferredoxin substrate. (A) Overlaid size exclusion chromatograms of FusB_CTD_ (blue), spinach ferredoxin (green) and a mixture of the two proteins (orange) with the arrow pointing at the ferredoxin-FusB_CTD_ complex. (B) ITC binding isotherm of 600 μM Fer_Sp_ titrated into 70 μM FusB_CTD_. The calculated Kd for the Fer_Sp_-FusB_CTD_ complex is 8.7(±3.5) μM (n=3). (C) Ferredoxin import assay showing the level of potato ferredoxin uptake by *ΔfusB* cells complemented with plasmids encoding either wild type (WT) or FusB_CTD_ R176K/R177K (2K).

Inspection of the amino acid sequence of FusB shows that the N-terminal portion of the predicted globular domain and the preceding linker contain a significant number of positively charged amino acids, which are absent from *Pc*LMG2410 and *E. coli* TonBs (Figure S7). Two such residues (Arg176, Arg177) are found in place of the highly conserved Gln-Pro-Gln residues, which form a part of the BtuB TonB-box binding motif (QPQYP) in *E. coli* TonB^20^. This arginine motif is located within a loop/linker region of TonBs, connecting the periplasmic-spanning and globular domains. Substitution of the two FusB_CTD_ arginine residues with lysines rendered a folded protein that did not comigrate with ferredoxin in gel filtration or interact with the substrate in ITC experiments (Figure S8). Similarly, *Pc*LMG2410 *ΔfusB* could not be complemented with a pFusB plasmid encoding the FusB R176K/R177K variant (Figure 4c). Therefore at least one of these two arginine residues appears to be critical for FusB-substrate interaction.

## Discussion

Our recent discovery that ferredoxin is imported into the periplasm of *P. carotovorum* revealed an unprecedented example of protein uptake for nutrient acquisition in Gramnegative bacteria^11^. In this work, we define key aspects of the mechanism of ferredoxin transport across the outer membrane. In a recent report, it was hypothesised that the M16 protease FusC acts as a periplasmic anchor that facilitates ferredoxin uptake by means of Brownian-ratchet mechanism^15^. However, the data presented here are inconsistent with this model, showing that ferredoxin import is independent of FusC. Instead, ferredoxin uptake requires energy transduction from the PMF and the TonB-like protein FusB. Therefore, the mechanism of ferredoxin import shares some similarity with the mechanism of import of widely studied substrates of TBDRs, such iron siderophores and vitamin B_12_^4,21^. For these substrates, according to the currently accepted models of TonB-dependent transport, the major role of TonB is in the displacement or partial displacement of the plug domain from their specific TBDTs^6,10^.

Interestingly, in the case of FusA, both FusB and *Pc*TonB are able to interact with its N-terminal region and so both these proteins may be able to facilitate displacement of the FusA plug domain. However, deletion of the genes encoding the two TonB proteins showed that only FusB is essential for ferredoxin transport, demonstrating an additional role for FusB in this process that cannot be fulfilled by *Pc*TonB. Although the affinity of FusB_CTD_ for FusA_NTR_ is low (57 μM), comparable low affinity complexes have been described between TBDT TonB-binding peptides and TonB proteins. For example, the affinity reported for the TonB-like protein HasB interaction with 21-mer HasR N-terminal peptide is 25 μM^22^. Similarly weak interactions between TonB and TBDR TonB-box peptides have been reported for FhuA (36 μM)^23^ and BtuB (9.4 μM)^6^. However, complex formation between TonB and TonB-binding peptides is characterised by β-strand augmentation which is known to result in the formation of mechanically strong complexes^6^. Indeed, it has been demonstrated *in vitro* using atomic force microscopy that the TonB–BtuB Ton box complex is sufficiently mechanically robust to induce partial unfolding of the BtuB plug-domain, forming a channel through which the vitamin B_12_ substrate can translocate^6^.

The ability of FusB to form a complex with ferredoxin, which *Pc*TonB lacks, indicates that this additional role involves the direct interaction of FusB with the ferredoxin substrate and that this complex formation is essential for ferredoxin transport through the lumen of FusA. Consistent with this, we identified an arginine motif that is required for FusB-mediated ferredoxin uptake by *P. carotovorum* and formation of the FusB-ferredoxin complex. Our current model of Fus-mediated iron acquisition, whereby FusB fulfils two distinct roles, is schematically shown in Figure 5. In this model, binding of the substrate on the extracellular side of FusA releases the TonB-box into the periplasmic space, where it is captured by FusB. Due to the dimensions of the globular ferredoxin, which are similar to the lumen of its FusA TBDT^11^, ferredoxin is unlikely to be able to readily diffuse into the periplasm after removal of the FusA plug domain. We therefore hypothesise that the interaction of FusB with the substrate involves a further PMF dependent step required to pull the ferredoxin substrate through the lumen of FusA. This would involve the C-terminal domain of FusB, which is of comparable size to plant ferredoxins, to enter the lumen of FusA to contact ferredoxin on the cell surface. The FusB-ferredoxin complex can then be pulled into the periplasm, using the ExbBD complex and PMF, after which the substrate is processed by FusC. Although we present a model relying on a single FusB per import cycle, we cannot exclude the possibility that the removal of FusA plug and ferredoxin import would involve two separate FusB molecules.

**Figure 5.**
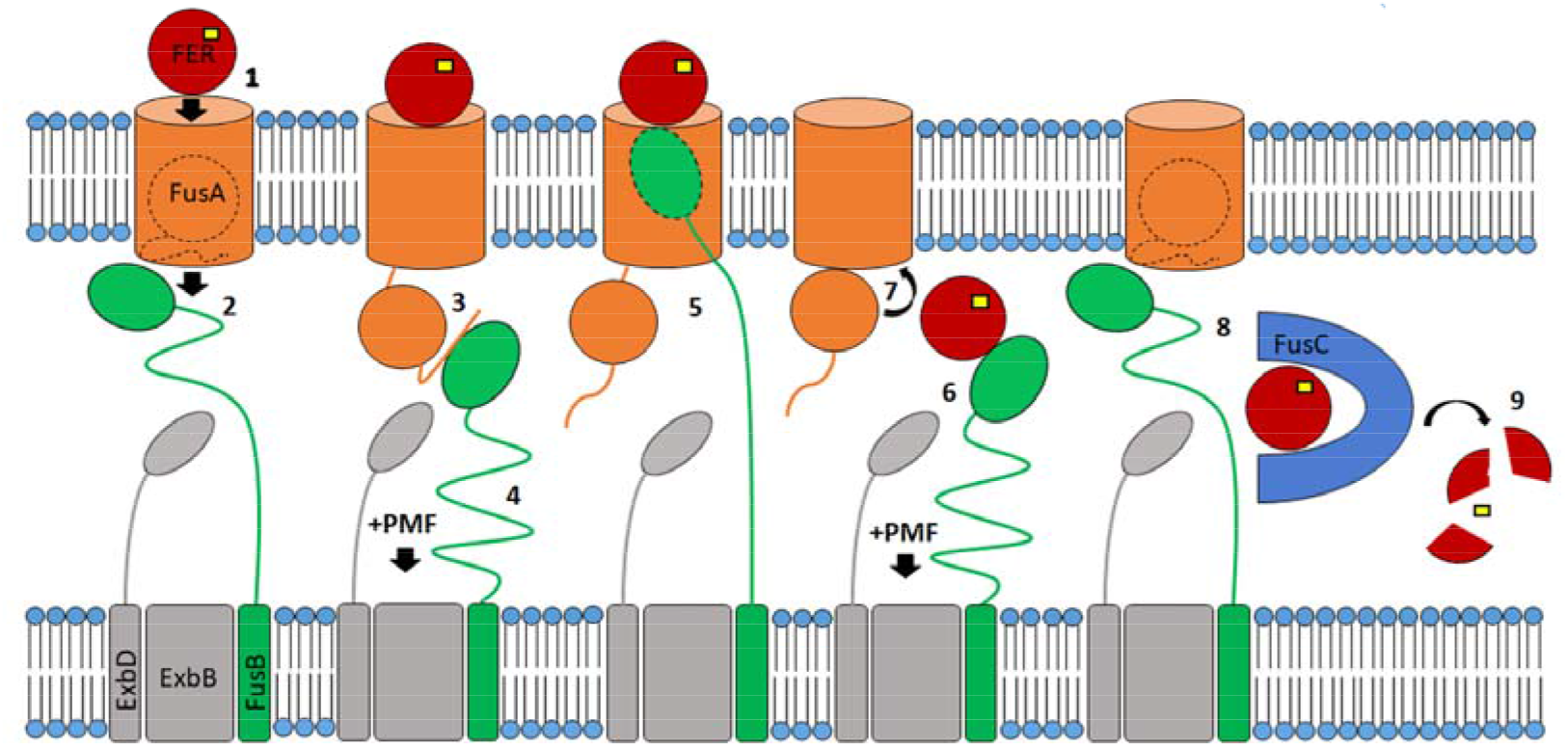
Proposed mechanism of FUS-mediated ferredoxin import mechanism. In the proposed mechanism FusB (green) fulfils two roles, firstly in displacement of the FusA plug domain and secondly in directly mediating ferredoxin translocation via the FusA lumen. Binding of ferredoxin (red) to FusA at the cell surface (1) causes release of the FusA TonB- box into the periplasm (2) where it is bound by FusB, which dislocates the plug domain (3) through energy transduced from the PMF via the ExbBD complex (4). FusB is then able to enter the lumen of FusA, bind ferredoxin (5) and through transduction of the PMF, translocate ferredoxin into the periplasm (6). The plug domain is then able to re-enter the FusA barrel (7) and return the FusA and FusB proteins to their resting states. Ferredoxin is then bound by FusC in the periplasm (8), which proteolytically cleaves the substrate releasing the iron-sulphur cluster (yellow) (9).

The occurrence of genes encoding multiple TonB-like proteins is a common feature of many Gram-negative bacteria^24^ and in some cases specific TonB proteins are required for the uptake of specific substrates, as for TonB2 of *Vibrio anguilarum* for anguibactin uptake^25^, while others exhibit some level of functional redundancy^26^. However, to our knowledge the Fus system represents the only substrate import system in which a TonB protein has been shown to directly interact with the substrate. This additional functionality displayed by FusB may reflect the nature of the ferredoxin substrate, which is atypically large in comparison to the well-studied TBDT siderophore substrates. In this respect, the uptake of ferredoxin is similar to the TonB-dependent uptake of the colicins and pyocins, which directly interact with TonB after threading their TonB box-containing intrinsically unstructured translocation domain (IUTD) though the lumen of their respective TBDT^10,27^. However, since plant ferredoxins are highly stable proteins that lack any kind of similar unstructured regions, our hypothesis is that in order to contact ferredoxin, FusB must enter the FusA lumen and contact FusA-bound substrate at the cell surface. This proposed mechanism also accounts for why the ferredoxin-containing bacteriocins do not require an IUTD that contains a TonB-box to cross the *P. carotovorum* outer membrane, with FusB able to directly contact their ferredoxin receptor-binding domains at the cell surface, thus enabling parasitisation of the Fus system^12,28^.

In summary, we describe a novel mechanism of ‘TonB-dependent’ nutrient uptake that requires a direct interaction between substrate and cognate TonB protein. The occurrence of multiple TonB proteins in many Gram-negative bacteria suggests that similar mechanisms may operate for atypical TBDT substrates.

## Material and Methods

### Bacterial strains and media

*E. coli* was grown in LB broth or plated on LB agar and grown at 37 °C. DH5α and BL21 (DE3) strains were used as host strains for cloning and for IPTG-induced protein expression, respectively. *P. carotovorum* was grown in LB broth or plated on LB agar at 30°C with the addition of the iron chelator 2,2’-bipyridine where specified. LB media and agar for culturing plasmid-complemented deletion strains always contained 100 μg ml^-1^ ampicillin.

### Generation of gene knockout strains and plasmids

The *fusA* (KAA3668913), *fusB* (KAA3668912), *fusC* (KAA3668914) and *tonB* (KAA3668374) sequences were determined from the genome sequence of *P. carotovorum* LMG2410 (GenBank: BioProject PRJNA543207)^29^. Genes in of *Pc*LMG2410 were deleted using the lambda red method as described previously^11,30^. The primers used for amplifying the kanamycin cassette from pKD4 template plasmid, gene sequences from genomic DNA and plasmid site-directed mutagenesis are listed in Table S1. The gene knockouts were confirmed by PCR and sequencing. Table S2 shows all the plasmids used in this study. To construct all plasmids, except pFusANTR-GFP, the respective genes were amplified from wild-type genomic DNA using primers that contained flanking regions with NdeI (forward) and XhoI (reverse) restriction enzyme sites. Purified PCR products were digested and ligated into NdeI/XhoI-digested pJ404, which carries ampicillin resistance. To generate pFusANTR-GFP the sequence encoding the N-terminal portion of FusA was amplified with primers containing XhoI (forward) and BamHI (reverse) restriction enzyme sites and the PCR products were inserted into XhoI/BamHI-digested pWaldo plasmid (Waldo et al 1999). The complementation plasmids were transformed into competent LMG2410 knockout strains by electroporation.

### Protein production and purification

FusC, FusB_CTD_, TonB_CTD_, FusA_NTR_-GFP and all ferredoxin proteins were overproduced in *E. coli* and purified as described previously^11,12^, except spinach ferredoxin (Fer_Sp_) which was purchased from Sigma. GFP alone used as a negative control in ITC was produced by cleavage of FusA_NTR_-GFP with TEV protease for 2 hours at RT, at 50:1 ratio. The resulting GFP-His_8_ was separated from residual TEV protease by size-exclusion chromatography and the removal of the N-terminal region of FusA was confirmed by SDS PAGE.

### Growth enhancement assays

Growth enhancement in the presence of ferredoxin was performed on solid media as previously described^11^. Briefly, 10 ml of 0.8% pre-cooled agar was supplemented with 50 μl of mid-log culture in LB media and poured onto an LB agar base containing 400 μM 2,2’-bipyridine (and 0.2 mM IPTG where specified). For plasmid based complementation, 100 μg/ml ampicillin was added to the LB media base. 4 μl of ferredoxin at specified concentration was spotted onto the solidified plate. For growth enhancement in liquid media bacteria were grown in M9 minimal media. 10 ml cultures were inoculated with 1 in 50 dilution of overnight LB cultures and upon reaching OD_600_=0.45 they were supplemented with 0.2 μM Fer_Ara_ and growth was monitored by measuring the OD_600_ for 6 hours.

### Ferredoxin internalisation and depletion assays

The time course of ferredoxin internalisation was initiated by supplementing LB cultures of wild type or *ΔfusC* cells (OD_600_=0.5) with 2,2’-bipyridine to a final concentration of 200 μM. Fer_Ara_ or Fer_Pot_ was added to a final concentration of 1 μM and the cultures were grown at 30°C with shaking over the specified time. At each time point a volume equivalent to 1 ml cell suspension at OD_600_= 0.5 was removed, the cells were spun down and treated with BugBuster (Merck) for soluble protein extraction. To determine if *ΔfusA* and *ΔfusB* strains can take up ferredoxin, LB cultures of WT and deletion strains at OD ~ 0.5 were supplemented with 200 μM 2,2’-bipyridine and 1 μM *Arabidopsis* or 5 μM potato ferredoxin. After 2 hours at 30°C with shaking 1 ml of cells was pelleted and soluble proteins were extracted using BugBuster (Merck). For internalisation experiments involving plasmid-complemented deletion strains, LB cultures were grown until OD_600_=0.4 was reached, whereupon 2,2’-bipyridine and potato ferredoxin were added. Δ*fusA*+pFusA and Δ*fusB*+pFusB cultures were split into two separate tubes, one of which was supplemented with IPTG to a final concentration of 0.2 mM. After 2 hours cells were harvested and subjected to BugBuster extraction as described above.

Depletion of *Arabidopsis* ferredoxin was monitored in 2 ml M9 minimal media cultures of WT, *ΔfusA* and *ΔfusB* strains over the course of 4 hours. Each culture, as well as 2 ml of uninoculated media (negative control), was supplemented 100 μM 2,2’-bipyridine and 1 μM Fer_Ara_. At each time point 50 μl of culture was removed from each tube and after pelleting the cells the supernatant was mixed with SDS loading dye.

The effect of dissipating PMF on ferredoxin uptake was determined using protonophore CCCP (Sigma), which was dissolved in DMSO to a final concentration of 10 mM. Mid-log cultures of wild type and *ΔfusC Pc*LMG2410 in M9 media were supplemented with 300 μM 2,2’-bipyridine and 2 ml of each culture was mixed with 18 μl DMSO and 2 μl CCCP stock (for 10 μM final CCCP concentration) or 20 μl CCCP stock (10 μM final CCCP concentration) or 20 μl DMSO for the “no CCCP” control. The cultures were mixed and incubated at room temperature for 10 min, after which wild type cultures were supplemented with 1 μM Fer_Pot_ and *ΔfusC* cultures with 0.2 μM Fer_Ara_. After 45 min incubation at 30°C with shaking, 1 ml of each culture was pelleted, washed with 0.5 ml PBS and subjected to BugBuster extraction.

### Ferredoxin cleavage assays

Cleavage reactions were performed in 10 mM Tris-HCl, pH 7.5, 50 mM NaCl, with 2 μM FusC and 250 μM ferredoxin at RT. At each time point 12 μl were removed and mixed with SDS loading dye. Proteins were resolved on a 16% SDS PAGE gel and visualised by Coomassie staining.

### Analytical size-exclusion chromatography

Proteins were concentrated to ~600 μM and 20 μl of TonB_CTD_ or the relevant construct of FusB_CTD_ was mixed with an equal volume of Fer_Sp_. The mixtures were then diluted with SEC buffer (20 mM Tris-HCl, pH 7.5, 150 mM NaCl) to 0.2 ml and loaded onto Superdex 75 10/300 column (GE Healthcare), pre-equilibrated in the same buffer. Each protein was also passed through the column individually for reference. The chromatograms were recorded at both 280 and 330 nm.

### Isothermal titration calorimetry

Experiments were performed on a MicroCal iTC_200_ instrument (Malvern) at 25°C in 10 mM Tris–HCl pH 7.5, 150 mM NaCl, with differential power set to 3. All proteins were dialysed against the ITC buffer overnight at 4°C except FusB_CTD_ and TonB_CTD_, which were passed through gel filtration in ITC buffer immediately before the experiment. Each type of titration was repeated at least once using different batches of purified proteins. 2 μl injections were used and titrations were continued until the signal represented heats of dilution. The magnitude of heats of dilution for each titrant was established in a separate experiment, where the titrant was injected into buffer. K_d_ values are expressed as the mean (±SEM).

## Abbreviations

Ara: Arabidopsis
CCCP: carbonyl cyanide chlorophenylhydrazone
CTD: C-terminal domain
Fer: ferredoxin
GFP: green fluorescent protein
ITC: isothermal titration calorimetry
IUTD: intrinsically unstructured translocation domain
NTR: N-terminal region
Pc: *Pectobacterium carotovorum*
PMF: proton motive force
Pot: potato
IPTG: isopropyl β-D-1 thiogalactopyranoside
SEC: size exclusion chromatography
Sp: spinach
TBDT: TonB-dependent transporter.

## Supplementary Information

**Figure S1.**
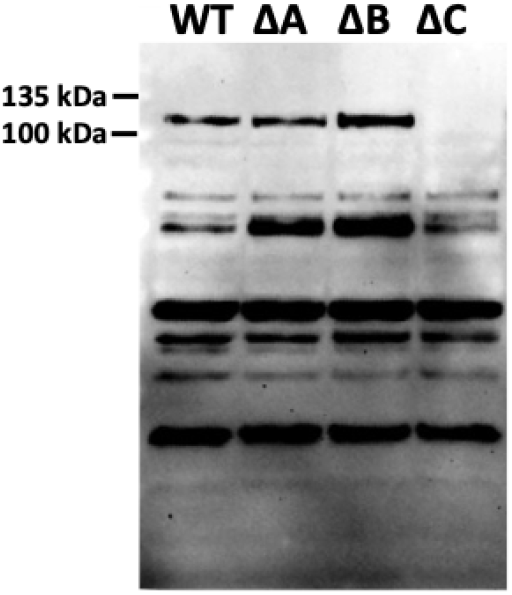
FusC is produced in *ΔfusA* and *ΔfusB* strains. Immunoblot showing the level of FusC (101 kDa) in the periplasmic fractions extracted from wild-type, *ΔfusA* (ΔA), *ΔfusB* (ΔB) and *ΔfusC* (ΔC) cells grown to mid-log phase in the presence of 2,2’-bipyridine.

**Figure S2.**
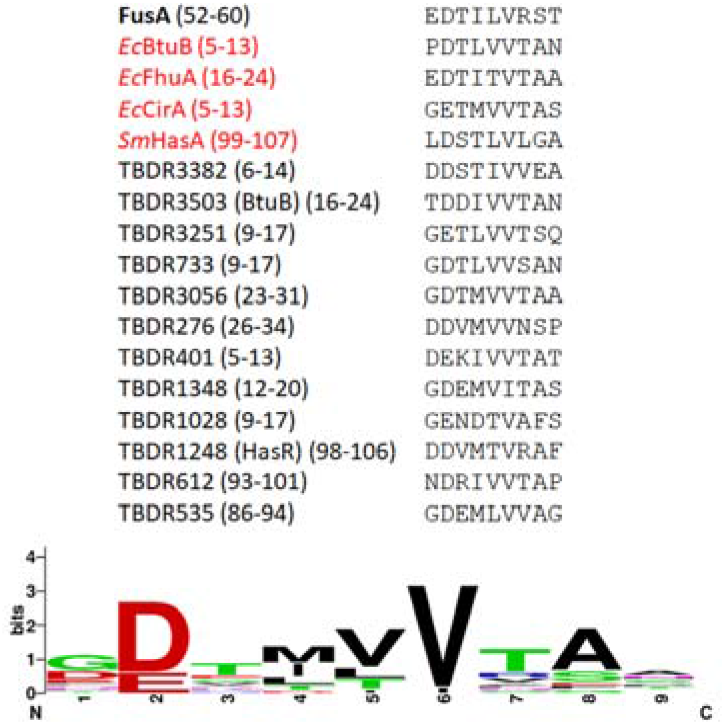
Sequence alignment of the putative TonB box regions of FusA against other Pc LMG2410 and *E. coli* TonB box regions. The numbers refer to the position of the predicted TonB box in the mature protein. Three Pc LMG2410 proteins were not included as their TonB boxes were not clearly identifiable. Sm refers to *Serratia marcescens,* Ec to *E. coli.* The consensus sequence shown below was generated using Weblogo.

**Figure S3.**
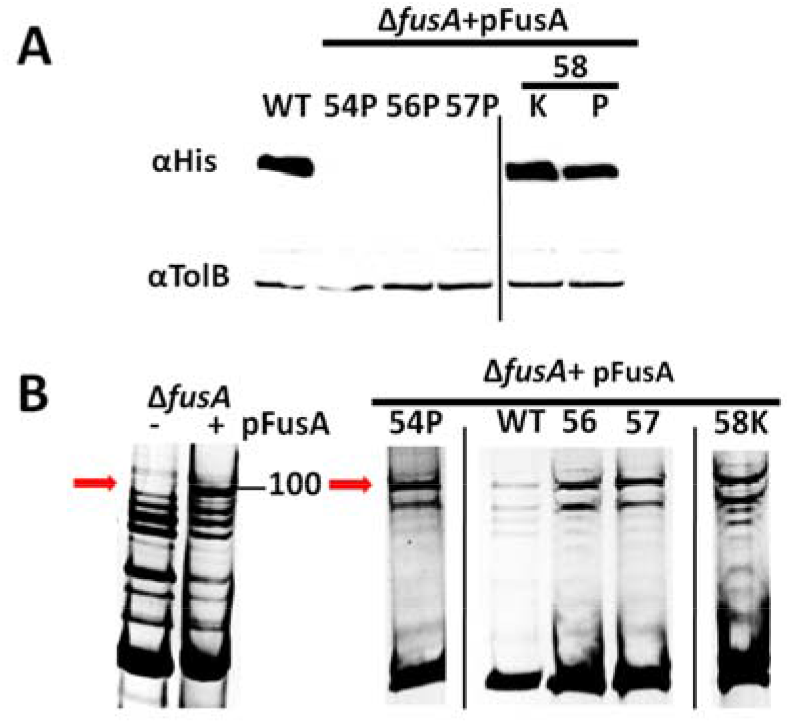
Mutations within the putative FusA TonB box affect ferredoxin transport. A - potato ferredoxin import assay of *ΔfusA* cells recomplemented with gene encoding either wild type or FusA variants. B - outer membrane extracts confirming the expression and appropriate localisation of wild type and mutant FusA.

**Figure S4.**
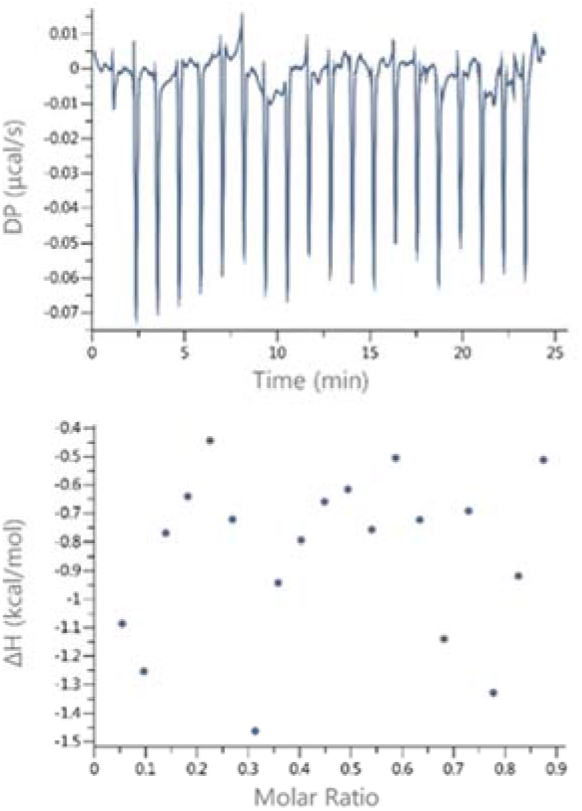
GFP does not interact with FusB_CTD_. Titration of 1.2 mM GFP, resulting from TEV mediated cleavage of FusA_NTR_-GFP, into 60 μM FusB_CTD_.

**Figure S5.**
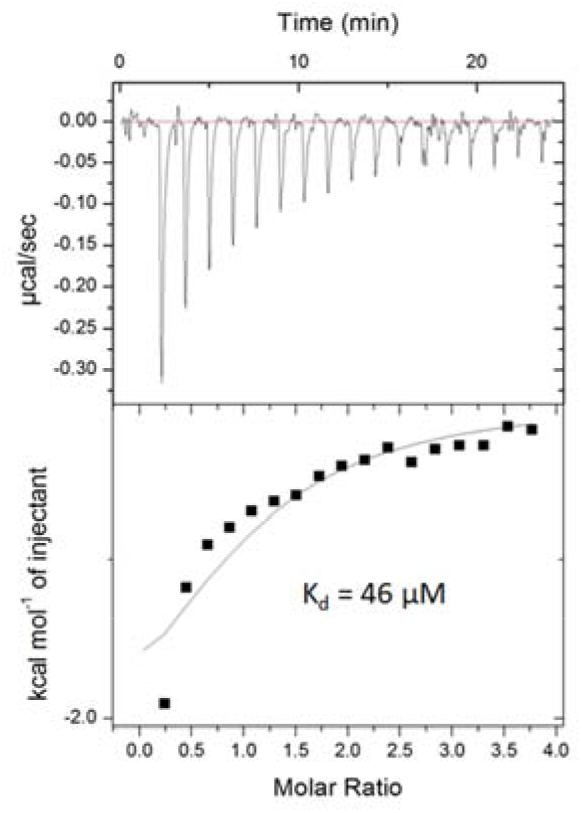
TonB_CTD_ forms a complex with the N-terminal domain of FusA. ITC was performed by titrating 1.2 mM FusA_NTR_-GFP into 45 μM TonB_CTD_.

**Figure S6.**
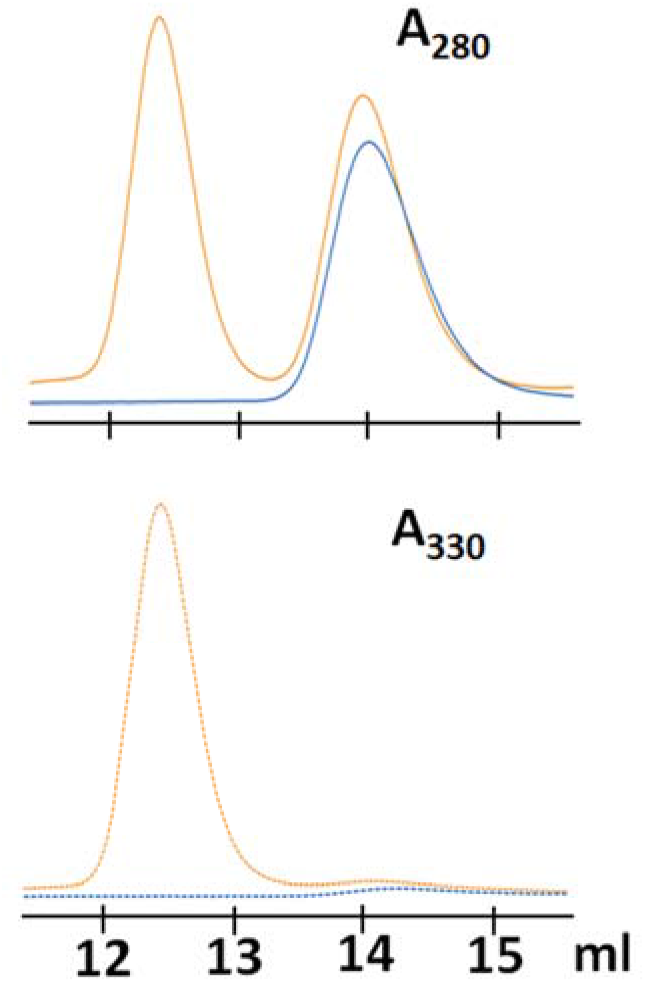
TonB_CTD_ does not form a complex with ferredoxin. Overlay of size-exclusion chromatograms showing TonB_CTD_ in the absence (blue trace) and in the presence of Fer_SP_ (orange trace).

**Figure S7.**
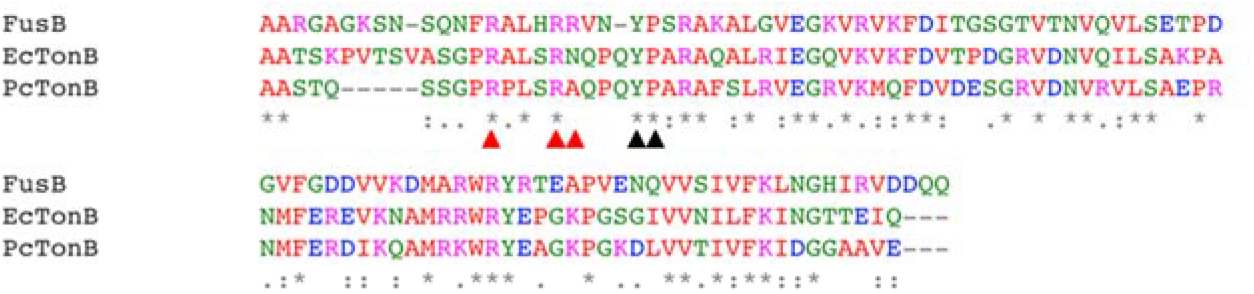
ClustalW sequence alignment of the C-terminal domains of FusB, *Pc*TonB and *Ec*TonB. Alignments start at residue 211 of FusB, 158 of *Pc*TonB and 140 of *Ec*TonB. Black arrows indicate the highly conserved Tyr and Pro residues, red ones show arginines in FusB that are present in the first loop of the C-terminal domain.

**Figure S8.**
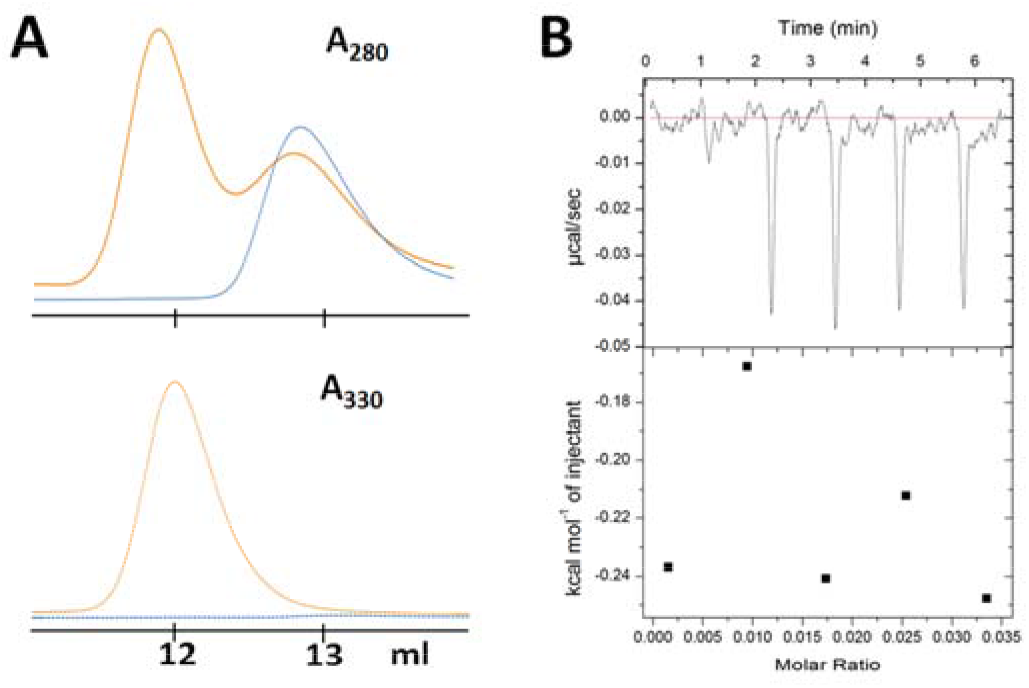
Substitution of two arginine residues in FusB_CTD_ with two lysines precludes interaction with ferredoxin molecule. A - size exclusion chromatograms at 280 and 330nm of FusB_CTD_2K alone (blue) or in the presence of spinach ferredoxin (orange) showing no complex formation. B - ITC titration of Fer_Sp_ into FusB_CTD_2K showing only heats of dilution.

**Table S1.**
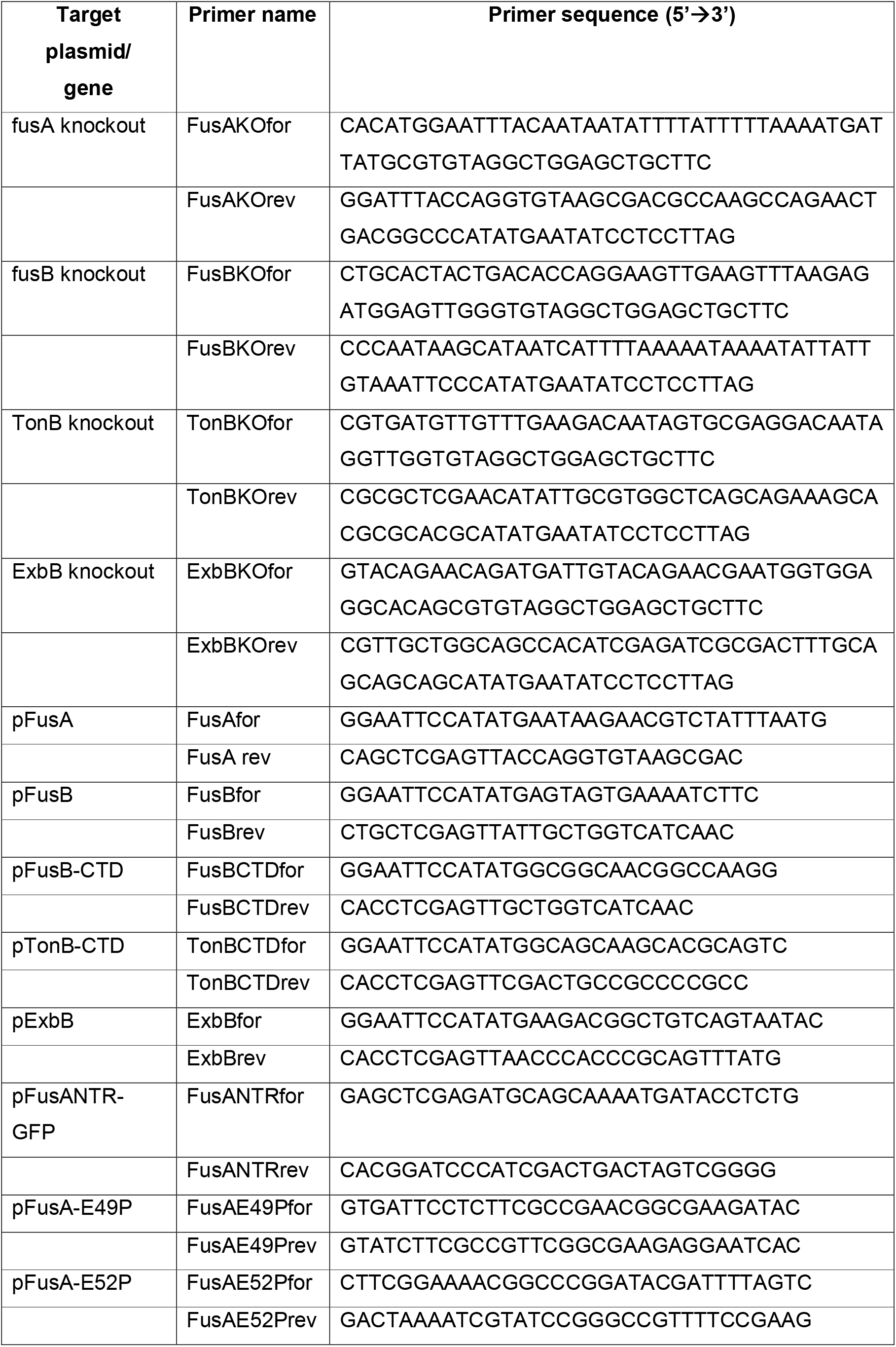
List of primers used in this study.

**Table S2.**
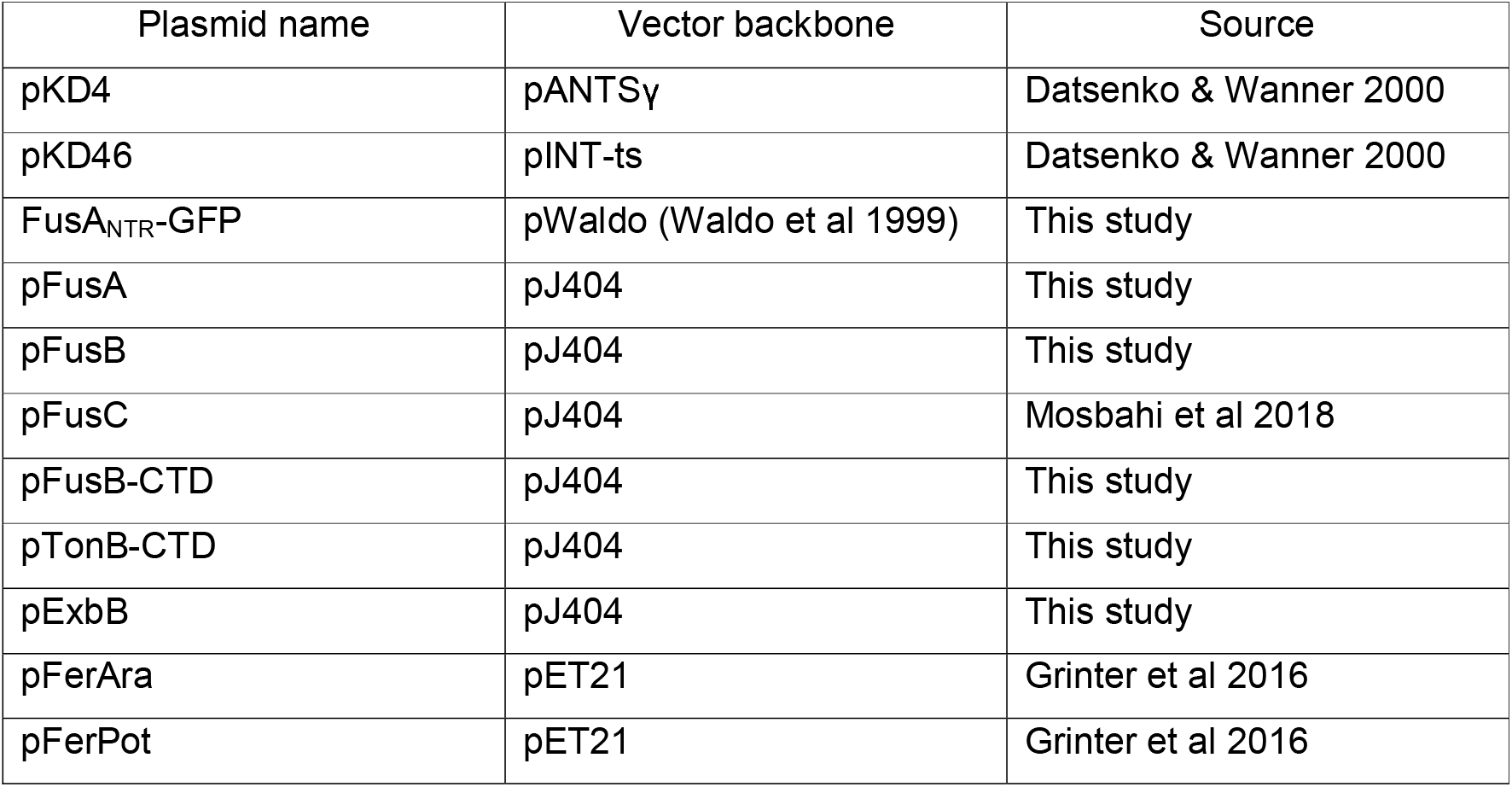
List of plasmids used in this study

